# An engineered, orthogonal auxin analog/_*At*_TIR1(F79G) pairing improves both specificity and efficacy of the auxin degradation system in *Caenorhabditis elegans*

**DOI:** 10.1101/2021.08.06.455414

**Authors:** Kelly Hills-Muckey, Michael A. Q. Martinez, Natalia Stec, Shilpa Hebbar, Joanne Saldanha, Taylor N. Medwig-Kinney, Frances E. Q. Moore, Mariia Ivanova, Ana Morao, Jordan D. Ward, Eric G. Moss, Sevinc Ercan, Anna Y. Zinovyeva, David Q. Matus, Christopher M. Hammell

## Abstract

The auxin-inducible degradation system in *C. elegans* allows for spatial and temporal control of protein degradation via heterologous expression of a single *Arabidopsis thaliana* F-box protein, transport inhibitor response 1 (_*At*_TIR1). In this system, exogenous auxin (Indole-3-acetic acid; IAA) enhances the ability of _*At*_TIR1 to function as a substrate recognition component that adapts engineered degron-tagged proteins to the endogenous *C. elegans* E3 ubiquitin ligases complex (SKR-1/2-CUL-1-F-box (SCF)), targeting them for degradation by the proteosome. While this system has been employed to dissect the developmental functions of many *C. elegans* proteins, we have found that several auxin-inducible degron (AID)-tagged proteins are constitutively degraded by _*At*_TIR1 in the absence of auxin, leading to undesired loss-of-function phenotypes. In this manuscript, we adapt an orthogonal auxin-derivative/mutant _*At*_TIR1 pair (*C. elegans* AID version 2 (*C.e.*AIDv2)) that transforms the specificity of allosteric regulation of TIR1 from IAA to one that is dependent on an auxin derivative harboring a bulky aryl group (5-Ph-IAA). We find that a mutant _*At*_TIR1(F79G) allele that alters the ligand binding interface of TIR1 dramatically reduces ligand-independent degradation of multiple AID*-tagged proteins. In addition to solving the ectopic degradation problem for some AID targets, addition of 5-Ph-IAA to culture media of animals expressing _*At*_TIR1(F79G) leads to more penetrant loss-of-function phenotypes for AID*-tagged proteins than those elicited by the _*At*_TIR1-IAA pairing at similar auxin analog concentrations. The improved specificity and efficacy afforded by the mutant _*At*_TIR1(F79G) allele expands the utility of the AID system and broadens the number of proteins that can be effectively targeted with it.

**ARITCLE SUMMARY:** Implementation of the auxin induced degradation (AID) system has increased the power if the *C. elegans* model through its ability to rapidly degrade target proteins in the presence of the plant hormone auxin (IAA). The current *C.e*.AID system is limited in that a substantial level of target degradation occurs in the absence of ligand and full levels of target protein degradation require high levels of auxin inducer. In this manuscript, we modify the AID system to solve these problems.

## INTRODUCTION

Detailed analyses of developmental and other dynamic biological events, processes and mechanisms have been facilitated in part by advancements in techniques to precisely control the products of gene expression. In *Caenorhabditis elegans*, a variety of tools have been developed to examine the stage- and tissue-specific function of genes, including FLP-mediated recombination to control gene activation(Davis *et al*. 2008; Voutev and Hubbard 2008) and RNAi to degrade the RNA gene product(Qadota *et al*. 2007). While these techniques are indirect, limited by the stability of the target protein, and are prone to off-target effects, more modern approaches have leveraged the power of CRISPR genome editing to engineer gene products to harbor epitopes that enable proteins to be directly targeted for degradation.

The auxin-inducible degradation system allows for the rapid and conditional degradation of target proteins in yeast, vertebrate cells, and *C. elegans* (Nishimura *et al*. 2009; Holland *et al*. 2012; Zhang *et al*. 2015). This two-component system relies on the heterologous expression of a plant-specific F-box protein, TIR1, that binds the plant hormone auxin (Ruegger *et al*. 1998; Gray *et al*. 1999) and an encoded 44-amino acid minimal degron sequence (auxin-inducible degron (AID*)) derived from the *Arabidopsis thaliana* IAA17 protein(Morawska and Ulrich 2013). The tractability and specificity of this system to provide exquisite spatiotemporal control of target protein levels relies on two key features of the TIR1/AID*-tagged protein interaction. First, stable association of TIR1 with proteins harboring an AID* sequence is allosterically regulated by auxin (indole-3-acetic acid, or IAA)(Tan *et al*. 2007). Second, *A. thaliana* TIR1 can interact with endogenous members of the Skp1 and Cullin family of proteins to form a functional, ligand-gated ubiquitin E3 ligase that targets AID*-tagged proteins to the 26S proteosome(Nishimura *et al*. 2009; Kanke *et al*. 2011; Holland *et al*. 2012; Zhang *et al*. 2015).

The AID system has proven to be extremely powerful in the *C. elegans* model where targeted inactivation of proteins can complement the already robust genetics of this system. The utility in this model comes from the application of many tissue-specific drivers that have been employed to drive TIR1 expression in various cell types(Ashley *et al*. 2021). Three issues have limited the applicability of this system for general use. First, the required dose of natural auxin (IAA) typically required for efficient degradation of target proteins is relatively high (1mM) and can elicit defined biological responses in *C. elegans* independent of _*At*_TIR1 expression (Bhoi *et al*. 2021; Loose and Ghazi 2021). The current system also exhibits a substantial amount of activity against a simple, heterologous AID*::GFP reporter even in the absence of added IAA (Martinez *et al*. 2020). Whether ectopic _*At*_TIR1-mediated degradation is a general issue of the AID system or a reflection of latent properties of individual AID*-tagged target proteins in *C. elegans* is *unknown*. Alternatively, endogenously- or environmentally-derived partially activating ligands may be present in *C. elegans* and these metabolites may trigger aberrant AID-target degradation in the presence of _*At*_TIR1. Finally, some AID-targets are inefficiently degraded and fail to generate strong loss-of function phenotypes (Patel and Hobert 2017; Serrano-Saiz *et al*. 2018; Duong *et al*. 2020).

In this manuscript, we aimed to improve the *C. elegans* AID system by altering the ligand binding specificity of TIR1. Using a previously established TIR1 variant that can target degradation in *A. thaliana* using a synthetic, modified auxin/IAA ortholog (Uchida *et al*. 2018), we demonstrate that the _*At*_TIR1(F79G) variant combined with a 5-Ph-IAA auxin analog also functions in the *C. elegans* model to degrade AID*-tagged proteins. Importantly, substitution of this single amino acid in _*At*_TIR1 alleviates the ligand-independent activity of this protein for multiple, biologically relevant AID*-tagged fusion proteins. Finally, we demonstrate that the _*At*_TIR1(F79G)/5-Ph-IAA (*C*.*e*.AIDv2 System) also exhibits elevated activity, enabling strong loss-of-function phenotypes to be elicited with low levels of exogenously added ligand. Because the AIDv2 system efficiently targets existing AID*-tagged target genes, we suggest that this modification functions as an improved experimental platform to dissect spatial and temporal gene functions.

## MATERIALS AND METHODS

### *C. elegans* maintenance and genetics

*C. elegans* strains were maintained on standard media at 20°C and fed *E. coli* OP50(Brenner 1974). Some strains were provided by the CGC, which is funded by NIH Office of Research Infrastructure Programs (P40 OD010440). A complete list of strains outlined in this manuscript can be found in Supplemental Table 1. 400mM IAA (Sigma; Product #I3750) and 100mM 5-Ph-IAA (Bioacademia; Product #30-003-10) stocks were made in 95% ethanol diluted into NGM media at the indicated concentration.

### CRISPR Editing

#### cshIs140(TIR1(F79G))

The single copy *rps-28pro::*_*At*_TIR1*::T2A::mCherry::his-11* (*cshIs128*) was integrated using standard CRISPR-mediated genomic editing to the ttTi5606 landing site following standard protocols(Dickinson *et al*. 2013). CRISPR editing of the *cshIs128* allele to generate the _*At*_TIR1(F79G) variant was accomplished using standard procedures (Paix *et al*. 2017). Briefly, recombinant nlsCas9 obtained from the University of California, Berkley MacroLab was used in conjunction with a recombinant sgRNA (Synthego, Menlo Park, California) with the following guide sequence (PAM sequence underlined): TTCCCTTGAGCTCGACGGAA<u>CGG</u> (Oligo #1; see oligo table). A single stranded, HPLC-purified repair template (Oligo #2; see oligo table) was used to edit the TIR1 coding sequence by homologous repair. The allele was verified by standard Sanger sequencing.

#### lin-28::AID*(ae157)

The AID*-tagged *lin-28* allele was constructed in a similar manner as above using a separate commercially available Cas9 protein (EnGen® Spy Cas9 NLS, M0646T) with gRNA targeting a genomic sequence in the 3’ coding sequence of *lin-28* (5’-ATATCATCGTCAGATGTAGT-3’). The sgRNA was synthesized using the Invitrogen™, MEGAshortscript™ T7 Transcription Kit (AM1354) and a single stranded transcription oligo #1 template (oligo #3; see oligo table). The repair templates were generated using to template oligos #4 and #5 (see oligo table). Repair templates #2 and #3 were mixed together, heated to 96 °C for 5 minutes, then placed on ice. The purpose was to create a “hybrid template” that is more efficient in the homology-directed repair(Dokshin *et al*. 2018).

#### ama-1::AID*::GFP(ers49)

The 1752bp repair template to generate the AID*::GFP C-terminally tagged *ama-1* allele was made by amplifying the AID*::GFP sequences from pLZ29(Zhang *et al*. 2015) using the oligos #6 and #7 (see oligo table). Injection mixtures using recombinant *S. pyogenes* Cas9 3NLS (10 μg/μL, IDT), crRNA (GATGAATTTGGATCATAAGT, 2 nmol, IDT, oligo #8; see oligo table), tracrRNA (IDT, cat# 1072532) dsDNA donors, pCFJ90 (co-injection marker pharynx mCherry was used at a final concentration of 3 ng/µL) were prepared as previously described(Dokshin *et al*. 2018).

#### hrpa-1::AID*(zen91)

To generate *hrpa-1::linker::AID*::TEV::FLAG* strain, N2 worms were injected with the CRISPR-Cas9 RNA-protein complex(Paix *et al*. 2017). The injection mix consisted of Alt-R Cas9 (IDT, cat# 1081058) along with *hrpa-1* crRNA (oligo #9; see oligo table), *dpy-10* crRNA (5’-GCUACCAUAGGCACCACGAG-3’)(Arribere *et al*. 2014), tracer RNA (IDT, cat# 1072532) and *hrpa-1::linker::AID*::TEV::FLAG* donor (oligo #10; see oligo table). Four independent alleles of *hrpa-1::linker::AID*::TEV::FLAG* were obtained and the tagged endogenous loci were sequenced. Two lines were chosen, outcrossed twice and assessed. One line was chosen for further experiments.

### Image acquisition

#### Confocal Microscopy

Images were acquired using a Hamamatsu Orca EM-CCD camera and a Borealis-modified Yokagawa CSU-10 spinning disk confocal microscope (Nobska Imaging, Inc.) with a Plan-APOCHROMAT x 100/1.4 or 40/1.4 oil DIC objective controlled by MetaMorph software (version: 7.8.12.0). Animals were anesthetized on 5% agarose pads containing 10mM sodium azide and secured with a coverslip. Imaging on the microfluidic device was performed on a Zeiss AXIO Observer.Z7 inverted microscope using a 40X glycerol immersion objective and DIC and GFP filters controlled by ZEN software (version 2.5). Images were captured using a Hamamatsu C11440 digital camera. For scoring plate level phenotypes, images were acquired using a Moticam CMOS (Motic) camera attached to a Zeiss dissecting microscope.

#### Wide-field Fluorescence microscopy

Images were acquired with a Zeiss Axio Observer microscope equipped with Nomarski and fluorescence optics as well as a Hamamatsu Orca Flash 4.0 FL Plus camera. An LED lamp emitting at 470 nm was used for fluorophore excitation. For single images, animals were immobilized on 2% agarose pads supplemented with 100mM Levamisole (Sigma).

### Image processing and analysis

All acquired images were processed using Fiji software (version: 2.0.0-rc-69/1.52p)(Schindelin *et al*. 2012). To quantify VPC-specific AID*::GFP or AMA-1::AID*::GFP expression levels, images were captured at the P6.p 1-cell stage (early L3 stage). Images of AID*::GFP animals were obtained at time points 0, 30, 60, 90, and 120 minutes in the absence or presence of auxin, and images of AMA-1::AID*::GFP animals were obtained 2 hours post-treatment. Expression levels were quantified by measuring the mean fluorescence intensity (MFI) of VPCs subtracted by the MFI of a background region in the image to account for camera noise. Cells were outlined using the freehand selection or wand (tracing) tool in Fiji. Kinetic data from AID*::GFP animals were normalized by dividing the MFI in treated or untreated animals at time points 30, 60, 90, and 120 minutes by the average MFI in untreated animals at 0 minutes.

### Embryonic viability and brood size measurements

Brood size measurements were carried out at 20°C and 25°C for each indicated strain using standard protocols (Tissenbaum and Ruvkun 1998). Briefly, single, late L4-stage animals of each genotype were transferred to a new plate every 24 hours for a total of four days. Plates were scored 24 and 48 hours after each transfer for the number of eggs and number of viable, hatched worms. The embryonic viability from each animal was established by calculating the percentage of unhatched eggs after 48 hours for each associated plate in the experiment. The total brood size was determined for individual worms by summing the number of viable offspring of each derived plate. In all cases the stated brood size is the average of >15 worms for each genotype. For *hrpa-1::linker::AID*::TEV::FLAG* brood measurements, embryos of each genotype were plated onto control, IAA, or 5-ph IAA plates and cultured until L4 stage. Single, late L4-stage animals of each genotype were transferred to a new plate. Larval arrest from each animal was established by calculating the percentage of arrested larvae that failed to grow after 24 hours.

### Motility assays

Motility assays are modified from a previously published protocol (Tsalik *et al*. 2003). Briefly, 5 animals (grown in the indicated conditions) were transferred to a standard NGM plate (± auxin analog) to a region in the center of a set of concentric rings (4mm, 6mm, 8mm, 10mm)(see Template 1). Animals were allowed to freely move for the indicated time and the proportion of animals that left the first ring within the annotated time were recorded. Animals were scored positive even if they left the ring and returned within the boundary.

### Western blotting

40-50 worms were picked directly into the 2x Laemmli protein loading buffer and boiled at 95°C for 5 minutes. HRPA-1 was detected using the Monoclonal ANTI-FLAG M2-Peroxidase (HRP) antibody (A8592-2MG) from Sigma-Aldrich at a 1:500 dilution. Mouse anti-tubulin antibody (Sigma-Aldrich) was used at a 1:5000 dilution to detect tubulin as a loading control.

### Graph Plots and Statistical Analysis

Plots and diagrams were generated using GraphPad Prism v9 (GraphPad Software, San Diego, Ca). Statistical significance was determined using a two-tailed un-paired Student’s t-test. P<0.05 was considered statistically significant. **** indicates P < 0.0001.

## Data availability

Strains in Supplementary Table S1 are available through the *Caenorhabditis Genetics* Center. Other strains and plasmids can be requested directly from the authors. The data that support the findings of this study are available upon reasonable request.

## RESULTS

### _*At*_TIR1(WT) triggers loss-of-function phenotypes of a LIN-28::AID* allele in the absence of exogenously-added ligand

During post-embryonic development, genes in the heterochronic pathway control the sequence of stage-specific cell divisions and cell fate specification (Rougvie and Moss 2013). A key feature of this pathway is that transitions from one stage-specific pattern of development to the next are controlled by the sharp temporal downregulation of protein coding genes (Rougvie and Moss 2013). The *lin-28* gene encodes a highly conserved RNA-binding protein that functions to modulate miRNA processing, turnover, and activity; *lin-28* is acutely down-regulated after the first larval stage (Moss *et al*. 1997; Lehrbach *et al*. 2009; Van Wynsberghe *et al*. 2011). *lin-28* null mutants, *lin-28(ga94)*, initiate post-embryonic development normally but then skip L2-stage patterns of cell division and immediately execute L3 patterns of development after the L1 molt (Fig. 1A) (Moss *et al*. 1997). Consequently, *lin-28(0)* mutants exhibit vulval morphology defects (protruding vulva; *pvl*), fail to proliferate their lateral seam cells, and precociously deposit adult-specific alae structures one stage earlier than wild-type animals (Figure 1B, Table 1).

**Figure 1.**
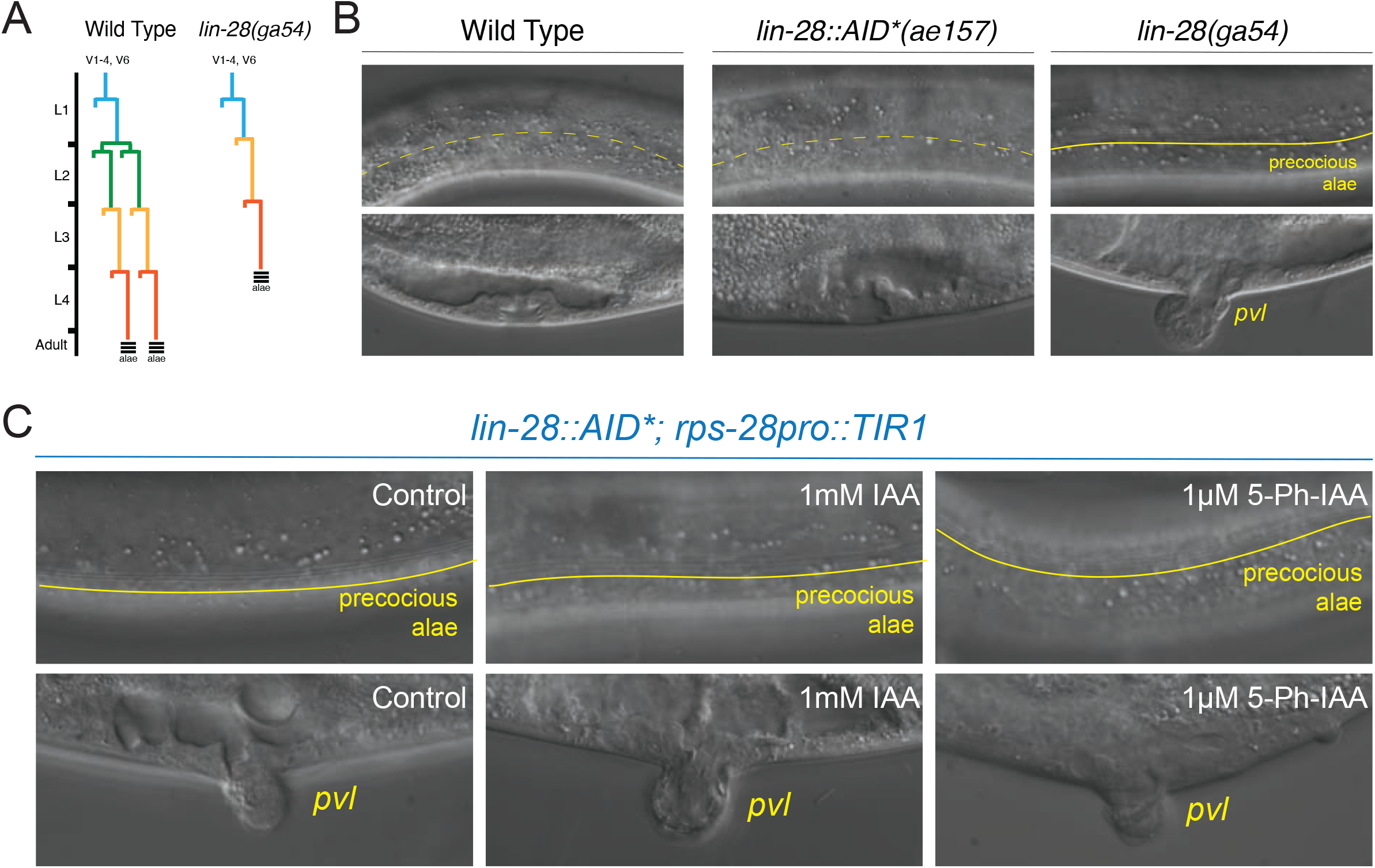
_At_TIR1(WT) induces a *lin-28* loss-of-function phenotype in the absence of exogenously added auxin ligand. **(A)** The lateral seam cells of wild-type *C. elegans* larva exhibit a stereotyped cell division program that generates additional seam cell in the L2 stage. Mutations in the heterochronic gene *lin-28* result in an altered seam cell division pattern due to the skipping of L2 stage-specific developmental programs. In addition, the lateral seam cells *lin-28(0)* mutants precociously exit the cell cycle and inappropriately deposit alae, adult-specific cuticle structures, during the L3-L4 molt. **(B)** In contrast to the normal skin and vulval developmental programs of wild-type animals and animals that harbor an AID-tagged *lin-28* allele, *lin-28(0)* mutants exhibit precocious alae at the L4 stage and a protruding vulval phenotype (pvl). **(C)** Combining the AID-tagged *lin-28* allele with a ubiquitously expressed *AtTIR1(WT)* allele results in strong heterochronic phenotypes in the absence of additional auxin. Dashed yellow lines indicate the absence of adult stage-specific alae structures whereas a solid line demarcates the presence of adult stage-specific alae structures. (See Table 1 for details).

**Table 1.** Measurement of TIR1- and _*At*_TIR1(F79G)-dependent heterochronic phenotypes in strains harboring an AID-tagged *lin-28* allele. A comparison and quantification of adult alae, seam cell numbers and percent of animals that exhibit protruding vulva phenotypes for animals expressing an AID-tagged *lin-28* allele.

| # | Strain | Genotype<sup>a,b</sup> | Experimental Condition<sup>c</sup> | % Animals L4 alae<sup>d</sup> | n = | % Animals Adult alae | n = | Ave. seam number<sup>d</sup> | Seam cell range | n = | % pvl animals | n = |
| --- | --- | --- | --- | --- | --- | --- | --- | --- | --- | --- | --- | --- |
| 1 | RG773 | Wild Type | - | 0 | 20 | 100 | 20 | 15.8 | 15---17 | 22 | 0 | 100 |
| 2 | HML1023 | *lin-28(ga54)I* | - | 85 | 20 | 100 | 15 | 11.8 | 10---12 | 27 | 100 | 100 |
| 3 | ME462 | *lin-28::AID(ae157) I* | - | 0 | 20 | 100 | 20 | 16 | 15---17 | 26 | 0 | 100 |
| 4 | HML1030 | *lin-28::AID(ae157) I;*<br>*cshIs128[rps-*<br>*28pro::TIR1_P2A_mCherryHis-11] II* | -<br>1mm auxin<br>1uM 5-Ph-IAA | 86<br>90<br>85 | 21<br>20<br>20 | 100<br>100<br>100 | 20<br>20<br>20 | 12.68<br>11.7<br>12.24 | 12---16<br>10---12<br>11---12 | 25<br>26<br>25 | 68<br>85<br>100 | 100<br>100<br>100 |
| 5 | HML1024 | *lin-28::AID(ae157) I;*<br>*cshIs140[rps-*<br>*28pro::TIR1(F79G)_P2A_mCherryHis-11] II* | -<br>1mm auxin<br>1uM 5-Ph-IAA | 0<br>0<br>91 | 20<br>20<br>23 | 100<br>100<br>100 | 20<br>20<br>20 | 16<br>15.8<br>11.7 | 16---17<br>15---16<br>11---12 | 42<br>31<br>23 | 0<br>0<br>100 | 100<br>100<br>100 |

<sup>a</sup>Animals contain *wIS78 V [ajm-1::GFP and scm::GFP]* which were used to visualize adherence junctions and lateral seam cells.
<sup>b</sup>L-the *ae157* allele of *lin-28* harbors an AID tag fused in frame to the *lin-28* coding sequence generating a carboxy-terminal tagged allele.
<sup>c</sup>L-4-staged Po animals were plated onto indicated plates and F1 progeny of the indicated stage were scored for
<sup>d</sup>Presence and quality of cuticular alae structures were assayed by Normarski DIC optics. Only one side of each animals was scored.
<sup>e</sup>Average seam cell numbers were scored by counting the number of *SCM::GFP(+)* cells on a single side of each animal.

To generate a *lin-28* allele whose expression can be modulated by auxin, we used CRISPR genome editing to insert a DNA fragment encoding the AID* tag at the 3’ end of the *lin-28* open reading frame. Examination of transgenic animals indicate that, in contrast to *lin-28(ga54)* animals, *lin-28::AID*(ae127)* animals do not exhibit heterochronic phenotypes, indicating that addition of the degron tag does not appreciably alter *lin-28* activity (Figure 2B and Table 1). We next crossed the *lin-28::AID*(ae127)* allele into a strain harboring a ubiquitously expressed *At*TIR1 (*cshIs128 [rps-28pro::TIR1::T2A::mCherry::his-11]*). Surprisingly, homozygous *lin-28::AID*(ae127)*; *chsIs128* animals exhibited highly penetrant heterochronic phenotypes (precocious vulval cell divisions, altered seam cell lineage (seam cell number and the precocious production of adult alae structures at the L4 stage) on normal NGM plates indicating that that the TIR1 activity, in the absence of auxin, results in a reduction of *lin-28* activity during development that phenocopies *lin-28(lf)* phenotypes.

**Figure 2.**
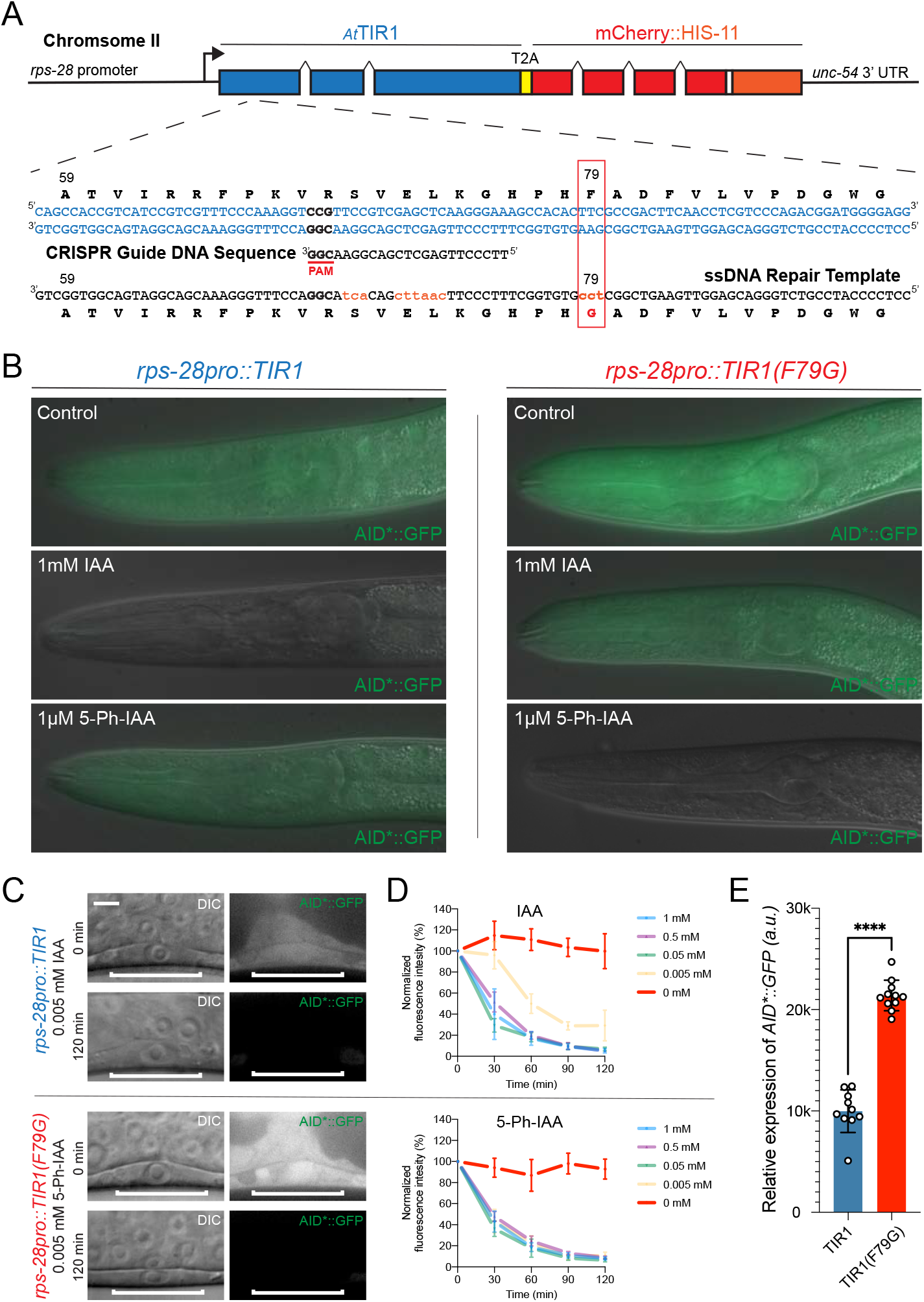
Mutation of phenylalanine 79 to glycine in the TIR1 protein switches the specificity of the auxin degradation system to one that is now responsive to 5-Ph-IAA. **(A)** Structure of *cshIs128* that encodes both a _*At*_TIR1(WT) protein and an autocatalytically-cleaved nuclear localized mCherry::HIS-11 reporter driven by a ubiquitously-expressed ribosomal protein (*rps-28*) promoter. Below the gene structure is the coding sequence of the region of _*At*_TIR1(WT) that was mutagenized via CRISPR and HDR using a co-injected ssDNA oligo. **(B)** Representative mid-larval staged animals expressing AID*::GFP and one of two indicated _*At*_TIR1 variants. Animals were grown continuously on untreated NGM plates or NGM plates including the indicated auxin analog at the listed concentration. **(C)** Micrographs of early L3 staged animals expressing AID*::GFP and one of the two _*At*_TIR1 variants before and 120 minutes after the addition of the auxin analog. **(D)** Rates of AID*::GFP degradation were determined by quantifying AID*::GFP in early L3 staged P6.p cells in animals co-expressing the indicated _*At*_TIR1 variant following auxin analog treatment. Data presented as the mean and SD (n ≥ 10 animals examined for each time point). **(E)** Quantification of the relative expression levels of the AID*::GFP reporter in P6.p cells of early L3 staged animals that were grown on control plates. Data presented as the median with SD (n ≥ 10 animals examined for each TIR1 transgene, and **** = p < 0.0001 by a Mann Whitney U-test).

### Generation, validation, and characterization of an orthogonal auxin and _*At*_*TIR1(F79G)* allele in *C. elegans*

Crystallographic studies indicate that IAA fills a hydrophobic cavity in between TIR1 and the AID/degron peptide sequence to form a stable trimeric complex (Dharmasiri *et al*. 2005; Kepinski and Leyser 2005; Tan *et al*. 2007). We hypothesized that the TIR1-dependent/auxin-independent degradation phenotypes of the *lin-28::AID\** allele could be caused either by a low-level interaction between the AID*-tag and TIR1 in the absence of auxin or by the inappropriate recognition of chemically-related endogenously produced ligand that may bind to the pocket of _*At*_TIR1. Uchida et al. (Uchida *et al*. 2018) have previously described a series of mutations of TIR1 that alter the auxin binding pocket to enable ligand-dependent target degradation using IAA derivatives that have been modified by aryl groups on the 5^th^ position of IAA. One of the engineered _*At*_TIR1 mutants they identified, _*At*_TIR1(F79G), failed to interact with AID*-tagged proteins in the presence of IAA but exhibited high binding specificity for AID*-tagged substrates in the presence of 5-phenyl-indole-3-acetic acid (5-Ph-IAA)(Uchida *et al*. 2018). We reasoned that the inappropriate activity of _*At*_TIR1, mediated by either of the above mechanisms, may be alleviated by similar alterations in the _*At*_TIR1-IAA binding pocket. A similar, orthogonal TIR1/auxin analog approach has been adapted for other systems using *Oryza sativa* TIR1 (*Os*TIR1) suggesting that this strategy is generally applicable (Yesbolatova *et al*. 2020).

To test the activity of a _*At*_*TIR1(F79G)* allele of TIR1 in *C. elegans*, we used CRISPR genome editing and homology-directed repair (HDR) to edit an existing, single copy TIR1 allele. Specifically, animals harboring a ubiquitously expressed *rps-28pro::TIR1* allele (*cshIs128*) and an *eft-3pro::AID*::GFP* transgene (*ieSi58*) were injected with a recombinant nlsCas9/guide ribonucleoprotein complex that is predicted to induce dsDNA breaks approximately 30bp upstream of the TIR1 F79 codon and a HRD repair oligo partially complementary to the TIR1 ORF overlapping with the TIR1 F79 codon (Figure 2). F1 progeny were then cloned onto NGM plates containing 1mM IAA to identify GFP(+) F2 progeny that fail to efficiently degrade the AID*::GFP reporter. Candidate GFP(+) F2 animals were then plated on NGM plates containing 100µM 5-Ph-IAA to identify clonal animals harboring a modified _*At*_TIR1 allele that is capable of degrading the AID*::GFP reporter only in the presence of the 5-Ph-modified auxin (Figure 2B). A single recombinant _*At*_*TIR1(F79G)* allele (*cshIs140*) was sequence-verified and used in further experiments.

We next compared the relative expression levels and degradation kinetics of the AID*::GFP reporter in animals expressing the two _*At*_TIR1 variants. Previous observations indicate that _*At*_TIR1(WT) reduces AID*::GFP expression in the absence of IAA (Martinez *et al*. 2020). We then compared the expression levels of the AID*::GFP reporter in animals expressing the two different TIR1 alleles. As shown in Figure 2C and E, AID*::GFP expression is significantly lower in the P6.p vulval precursor cells of animals expressing _*At*_TIR1(WT) when compared to the levels in similarly staged _*At*_TIR1(F79G)-expressing animals. Expression levels of AID*::GFP in _*At*_TIR1(F79G) animals is similar to that observed in WT animals (Figure S1). We next compared the efficiency of ligand-induced degradation mediated by these two _*At*_TIR1 alleles by quantifying the relative changes in AID*::GFP expression in the presence of various IAA or 5-Ph-IAA concentrations over time. We monitored AID*::GFP expression in single vulval precursor cells of stage-matched (early mid-L3 stage) animals that have been exposed to different concentrations of auxin orthologs that were incorporated into standard *C. elegans* solid culture media (Figure 2C and D). We found that elevated concentrations (≥ 0.05mM) of both auxin analogs led to the efficient reduction of AID*::GFP with greater than 50% of the target protein depleted in P6.p cells within the first 30 minutes of the time course (Figure 2D). At lower concentrations of 5-Ph-IAA (0.05mM), _*At*_TIR1(F79G) maintained similar AID*::GFP depletion kinetics suggesting that this level of activating ligand is still saturating in this system and that overall kinetics are dependent on _*At*_TIR1(F79G) expression levels. In contrast, the efficiency of the _*At*_TIR1(WT)-mediated degradation when normal IAA was reduced (Figure 2D). These results suggest that the _*At*_TIR1(F79G) (AIDv2) functions more efficiently with the 5-Ph-IAA ligand than the classical version of the AID system with unmodified IAA(Zhang *et al*. 2015).

### The AIDv2/TIR1(F79G) version of the AID system efficiently regulates the expression of dosage-sensitive AID* targets

Given that the _*At*_TIR1(F79G) allele of _*At*_TIR1 does not lead to a significant reduction of AID*::GFP expression in the absence of ligand, we next tested whether this allele could be used to efficiently modulate the activity of AID*-tagged genes that exhibit dosage-sensitive phenotypes in a ligand-specific manner. First, we crossed the _*At*_*TIR1(F79G)* allele into a strain harboring the *lin-28::AID\** degron-tagged allele. As opposed to the inappropriate *lin-28(lf)* phenotypes that are caused by _*At*_TIR1(WT), *lin-28::AID*; cshIs140 (*_*At*_*TIR1(F79G))* animals exhibit completely wild-type seam cell lineage (seam cell number) and do not exhibit precocious alae formation or a protruding vulval phenotype (Figure 3, Table 1). Importantly, treatment of *lin-28::AID*; cshIs140 (*_*At*_*TIR1(F79G))* parental animals with very low concentrations of 5-Ph-IAA (1µM) lead to highly penetrant heterochronic phenotypes for each developmental feature (seam cell number, alae, and vulval development) in F1 progeny indicating that the AIDv2 system can elicit fully penetrant *lin-28(lf)* phenotypes. These phenotypes were not elicited when animals were grown on media containing 1000-fold higher concentration of normal auxin (IAA), demonstrating the sensitivity and specificity of the TIR1(F79G) allele.

**Figure 3.**
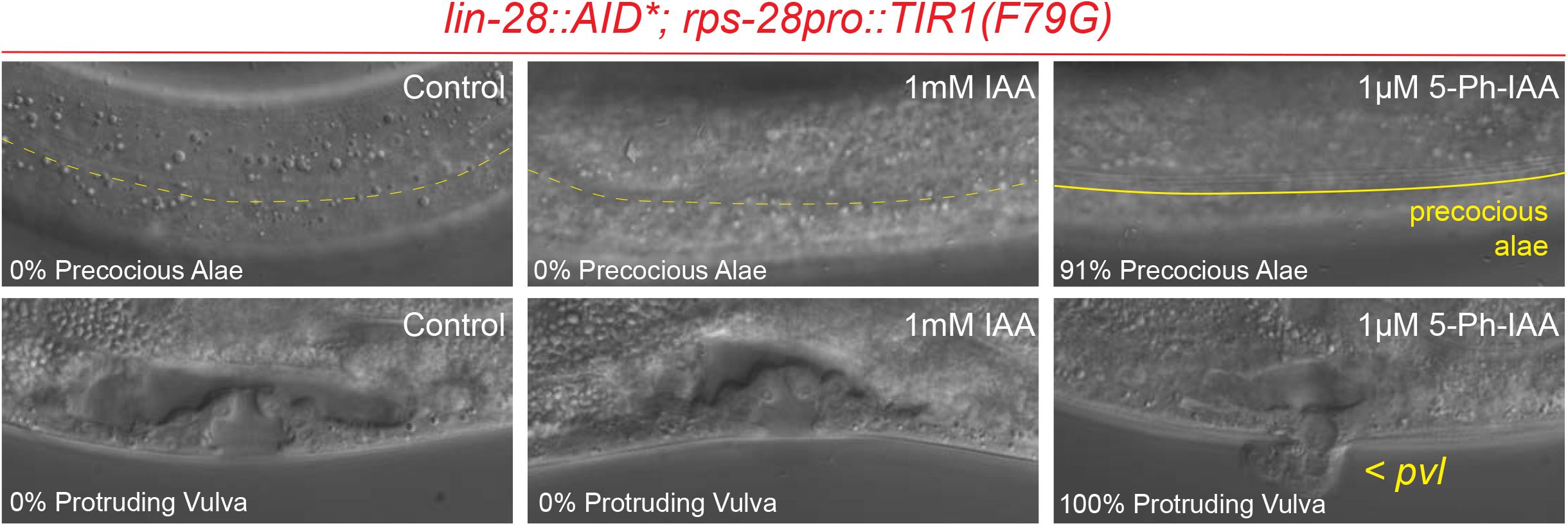
The _*At*_TIR1(F79G) variant of TIR1 allows for ligand-dependent regulation of LIN-28::AID* activity by 5-PhIAA. Representative images depict the cuticular and vulval development at the L4-stage of F1 animals that have developed on NGM plates containing the auxin analog at the indicated concertation. Dashed yellow lines demarcate the absence of adult stage-specific alae structures whereas a solid line demarcates the presence of adult stage-specific alae structures. Pvl = protruding vulva. (See Table 1 for details).

We have also previously tagged the large, catalytic subunit of the *C. elegans* pol II complex, *ama-1*, with the AID epitope with the idea that an auxin degradation system could be used to acutely inactivate transcription in a temporal and cell-type specific manner. This would bypass the limitations of prior approaches to reduce transcriptional output via chemical inhibitors or by cell-type-specific RNAi(Rogalski *et al*. 1988; Rogalski and Riddle 1988; Firnhaber and Hammarlund 2013). Transgenic animals homozygous for a C-terminally tagged *ama-1::AID*::GFP* allele, *ama-1(ers49)*, are indistinguishable from wild-type animals and exhibit normal development and brood sizes (Figure 4A). When the *ama-1::AID*::GFP* allele is combined with a ubiquitously expressed _*At*_TIR1(WT), the pace of overall animal development is dramatically slowed and there is a significant reduction (>8 fold) in the fecundity of animals compared to isogenic strains lacking _*At*_TIR1 expression (Figure 4A). Consistent with the assumption that these developmental phenotypes result from an inappropriate activity of _*At*_TIR1(WT), AMA-1::AID*::GFP expression is reduced by approximately 2.9-fold in P6.p vulval precursor cells (1-cell stage) of animals expressing ubiquitous _*At*_TIR1(WT) when compared to a strain only expressing the tagged pol II allele (Figure 4B and C).

**Figure 4.**
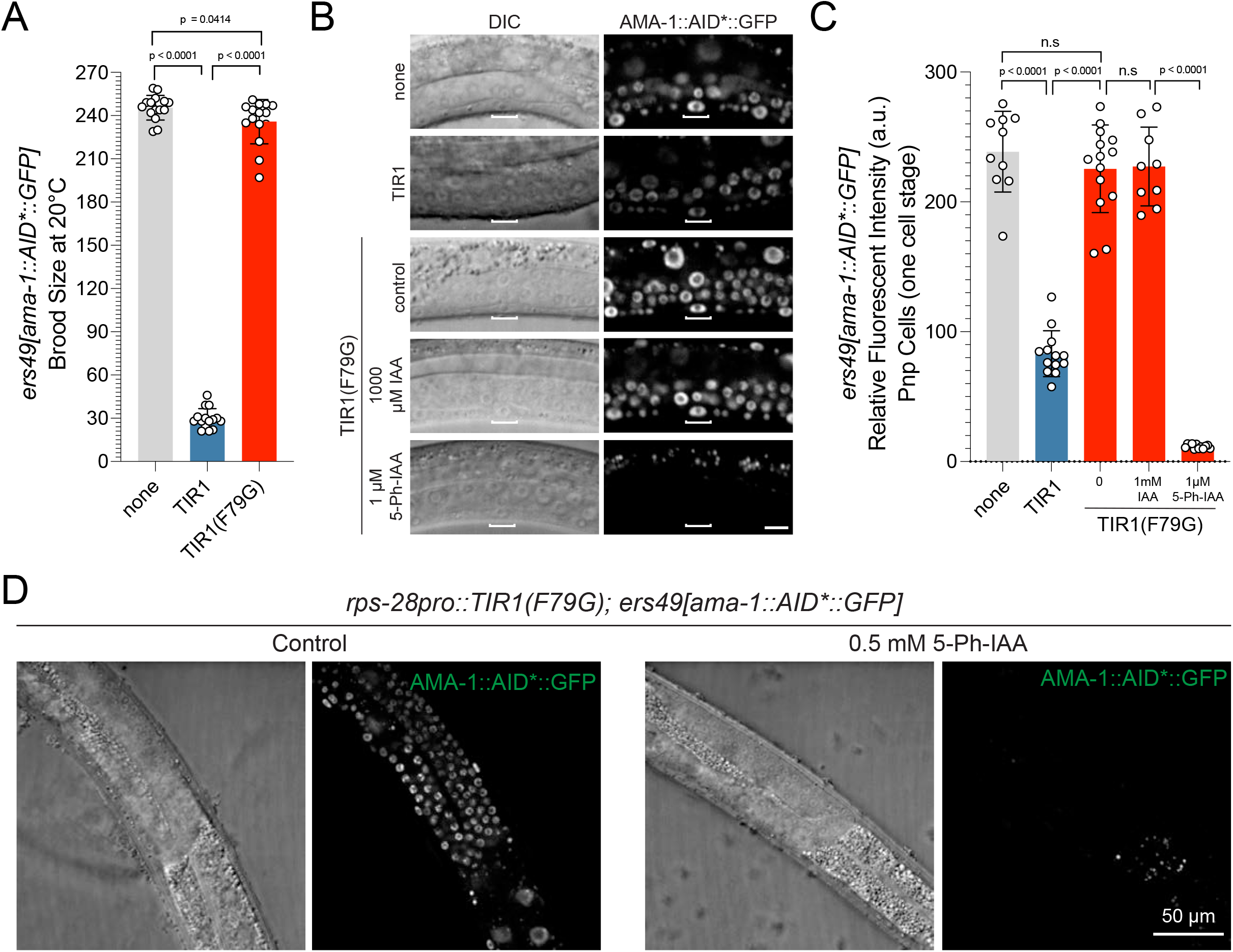
_*At*_TIR1(WT) allele of TIR1 generates ligand-independent loss-of-function phenotypes that are alleviated by the 5-Ph-IAA regulated _*At*_TIR1(F79G) variant. **(A)** An AID-tagged, endogenous allele of *ama-1, ama-1(ers49)*, exhibits a severe reduction in brood size when combined with _*At*_TIR1(WT) even in the absence of added auxin. In contrast, the _*At*_TIR1(F79G) variant does not alter the brood size of *ama-1(ers49)* animals (n = 15 for each genotype and significance was calculated using an unpaired t-test). **(B)** The expression of AMA-1::AID*::GFP is dramatically reduced in animals expressing _*At*_TIR1(WT) and not in animals expressing _*At*_TIR1(F79G). Shown are DIC and corresponding GFP images of VPCs (from early L3 stage animals at the P6.p 1-cell stage). Scale bar is 10 microns. **(C)** Quantification of AMA-1::AID*::GFP expression animals depicted in panel B. Data presented as the mean ± SD (n ≥ 10 animals examined for each, and P < 0.0001 by a Student’s t-test). **(D)** AMA-1::AID*::GFP is degraded in the germline by 5-Ph-IAA. Representative DIC and GFP images of L4 staged animals that have been transferred to control NGM plates or NGM plates containing 0.5 mM 5-Ph-IAA for 2 hours.

We next determined if the developmental phenotypes associated with the ectopic activity of _*At*_*TIR1(WT)* would also be alleviated by mutating the binding pocket of the _*At*_TIR1 ligand binding domain. When we crossed the *ama-1::AID*::GFP* allele into a strain expressing _*At*_*TIR1(F79G)*, we found that animals exhibited normal brood sizes, consistent with a lack of ligand-independent activity for this TIR1 variant (Figure 4A). In addition, AMA-1::AID*::GFP expression levels in _*At*_*TIR1(F79G)* expressing animals were similar to those observed in wild type (Figure 4B and C). Importantly, AMA-1::AID*::GFP expression is extinguished when animals are incubated with 5-Ph-IAA, but not a 1000-fold higher concentration of regular IAA (Figure 4B and C). AMA-1::AID*::GFP depletion results in a variety of pleiotropic, terminal phenotypes (sterility, slow growth, arrest, and lethality) that resemble phenotypes associated with *ama-1* mutations or animals that have been treated with alpha-amanitin, a pol II-specific enzymatic inhibitor (Rogalski *et al*. 1988; Rogalski and Riddle 1988; Bird and Riddle 1989). These results suggest that the _*At*_*TIR1(F79G)* allele of _*At*_TIR1 lacks appreciable ligand-independent activity on AID*-tagged substrates while maintaining the ability to program target degradation for multiple substrates.

### Exposure of *C. elegans* larva to high levels of IAA analogs elicits a transcriptional response

Previous experiments have demonstrated that indole (and likely indole derivatives) derived from commensal bacteria extends lifespan (Sonowal *et al*. 2017). Furthermore, recent reports utilizing the AID system have demonstrated that indole-3-acetic acid (IAA) at high physiological concentrations can also induce physiological responses in *C. elegans* that modulate a number of developmental and cellular activities. Specifically, continuous exposure of animals to auxin during development significantly extends the lifespan and can confer protection against endoplasmic reticulum (ER) stress (Bhoi *et al*. 2021; Loose and Ghazi 2021). Auxin antagonizes the negative effects of tunicamycin, a chemical inhibitor of N-linked glycosylation and a known inducer of ER stress, and this protective response requires the activity of the XBP-IRE-1 pathway of the Unfolded Protein Response (UPR)(Bhoi *et al*. 2021). These results collectively suggest that high levels of exogenous auxin may activate latent genetic programs that are optimized to ensure that animals can normally survive in diverse environments.

We discovered that auxin exposure induces the expression of multiple transcriptional reporters that are also induced in response to ER stress. These include transcriptional reporters for two glutathione S-transferases (*gst-4* and *gst-5*) and *gcs-1*, encoding a gamma glutamyl-cysteine synthetase, that are upregulated by SKN-1/Nrf, a transcription factor that orchestrates both oxidative and xenobiotic stress responses in *C. elegans* after exposure to tunicamycin (Figure 5A)(Papp *et al*. 2012; Glover-Cutter *et al*. 2013). The induction of this transcriptional response is not a generalized stress response as other transcriptional reporters of environmental stressors are not induced by auxin (e.g., *hsp-16::GFP*) (Figure 5A). Induction of *gcs-1pro::GFP* is rapid and peaks after 4-5 hours of 1mM auxin exposure (Figure 5B). This rapid and quantifiable response enabled us to both determine if 5-Ph-IAA also induces this regulatory pathway and then to identify the minimal dose of each auxin analog required to activate this *gcs-1pro::GFP* expression We found that both IAA and 5-ph-IAA induced *gcs-1pro::GFP* expression, but this induction was not significant over background until the auxin analog reached a concentration of greater than 0.5 mM (Figure 5C). Therefore, an optimal AID system should work at exogenously added auxin analog concentrations that, at a minimum, do not trigger this transcriptional response.

**Figure 5.**
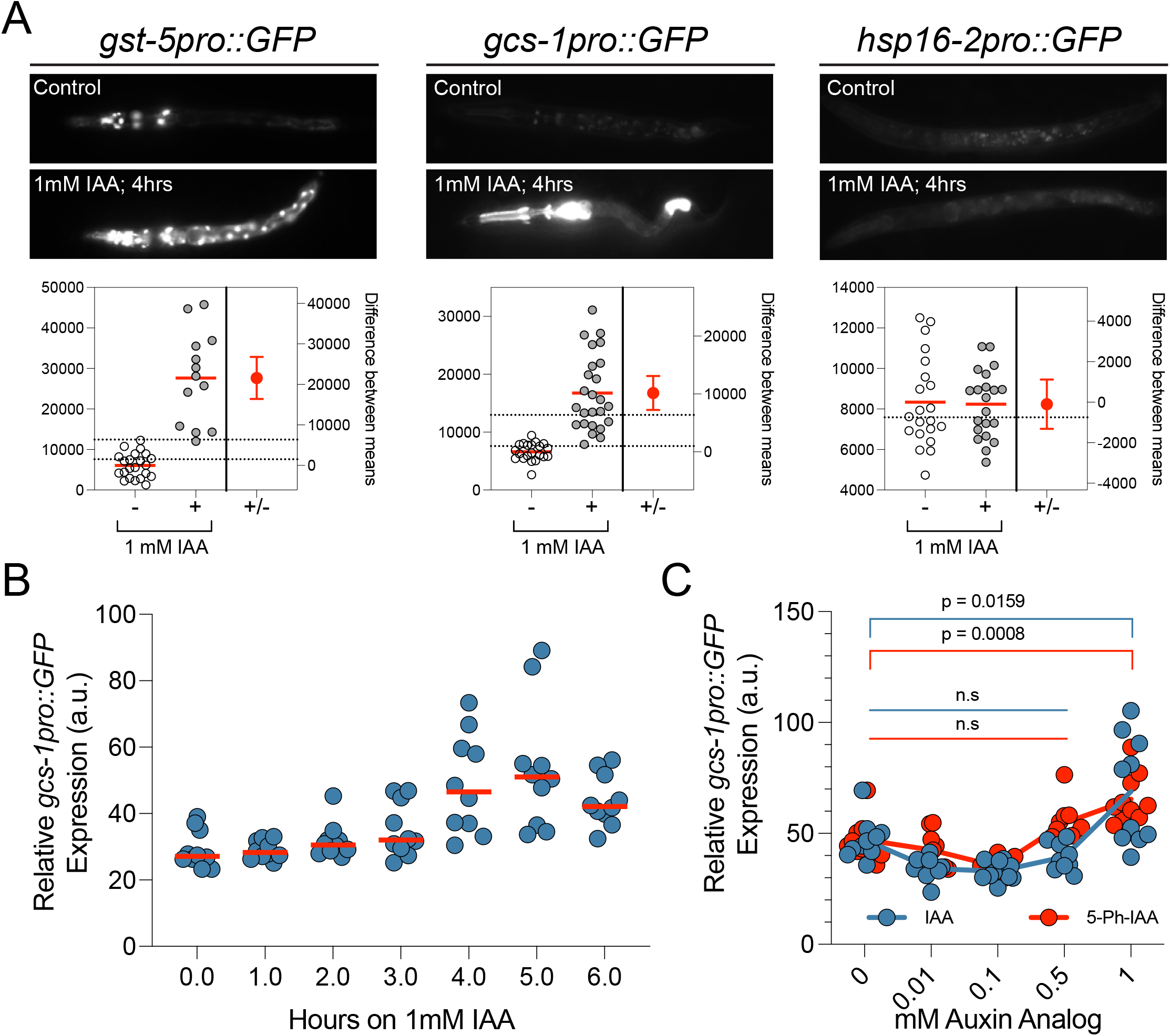
Elevated levels of auxin analogs induce stress-responsive gene expression. **(A)** Representative GFP images and estimation plots of the expression levels of each indicated stress-responsive reporter in animals that have been exposed to auxin/IAA for 4 hours. For quantification of GFP expression, the relative fluorescence of whole, individual animals was measured (n >13 for each reporter). **(B)** Transgenic *gcs-1pro::GFP* animals were exposed to 1mM IAA for the indicated times and relative fluorescence of individual animals was calculated as in Panel A. **(C)** Quantification of *gcs-1pro::GFP* expression levels in animals that have been incubated on plates containing the indicated concentrations of IAA or 5-Ph-IAA for 4 hours.

### The AIDv2 degradation system generates more penetrant loss-of-function phenotypes at lower IAA analog concentrations when compared to the classical TIR1(WT)-IAA pairing

Given that the typical high levels of IAA or 5-Ph-IAA trigger the ectopic expression of *gcs-1pro::GFP*, we aimed to determine if _*At*_TIR1(F79G) worked more efficiently than TIR1(WT) at lower auxin analog concentrations. Kinetic data comparing the two TIR1-auxin analog pairings in Figure 2C suggested that _*At*_TIR1(F79G) more efficiently depletes the expression of AID*::GFP than _*At*_TIR1(WT) in low auxin analog concentrations. To test this feature, we assayed both TIR1 variants for their ability to generate phenotypes on AID* targets that only display partially penetrant phenotypes with the AIDv1/TIR1(WT) system.

The *unc-3* gene, encoding the sole *C. elegans* ortholog of the Collier/Olf/Ebf (COE) family of TFs, functions as a neuronal terminal selector gene that is required to both establish and maintain distinct cholinergic motor neuron cell fates in the ventral nerve chord (Kerk *et al*. 2017; Feng *et al*. 2020; Li *et al*. 2020). Animals harboring non-functional alleles of *unc-3* display penetrant locomotion defects and exhibit a “coiler” phenotype where the posterior portions of the animal appear paralyzed (Figure 6A) (Brenner 1974). To determine if _*At*_TIR1(F79G) can elicit more penetrant phenotypes than _*At*_TIR1(WT) at similar auxin analog concentrations, we scored uncoordinated phenotypes of _*At*_TIR1(F79G) or *At*TIR1(WT) parental animals exposed to varied auxin analog concentrations using a modified radial motility assay (Tsalik *et al*. 2003) (Figure 6B). Specifically, late L4/adult F1 animals that had or had not been exposed to auxin analogs were transferred to a center point of a series of concentric circles of a marked NGM plate. Animals were then allowed to move freely for 2 minutes and the percentage of animals that moved greater than 2 mm from that center point were scored (Figure 6C). Using this assay, 100% of wild-type and ∼2% of *unc-3(e151)* animals moved > 2 mm within the recorded time course (Figure 6C). We found that treatment of _*At*_*TIR1(WT); unc-3::mNG::AID\** animals with IAA induced an uncoordinated phenotype but only at elevated IAA concentrations (approximately 45% of animals were motile enough to travel outside of the assay area (Figure 6C). In contrast, 5-Ph-IAA treatment of _*At*_TIR1(F79G) animals led to highly penetrant *unc* phenotypes in this assay (Figure 6C). Consistent with *unc-3* activity being continuously required for normal motility, _*At*_TIR1(F79G) animals treated after hatching also developed an *unc* phenotype by young adulthood, though the penetrance of these phenotypes was reduced when compared to continuous 5-Ph-IAA treatment (Figure S1). We do note that the expressivity of the *unc-3* phenotypes elicited by 5-Ph-IAA treatment in _*At*_*TIR1(F79G)*; *unc-3::GFP::AID\** animals, as measured by the proportion of animals that exhibiting a “coiler” phenotype, is lower than what is observed for strong *unc-3* loss-of-function alleles and are consistent with residual *unc-3* activity in 5-Ph-IAA treated animals.

**Figure 6.**
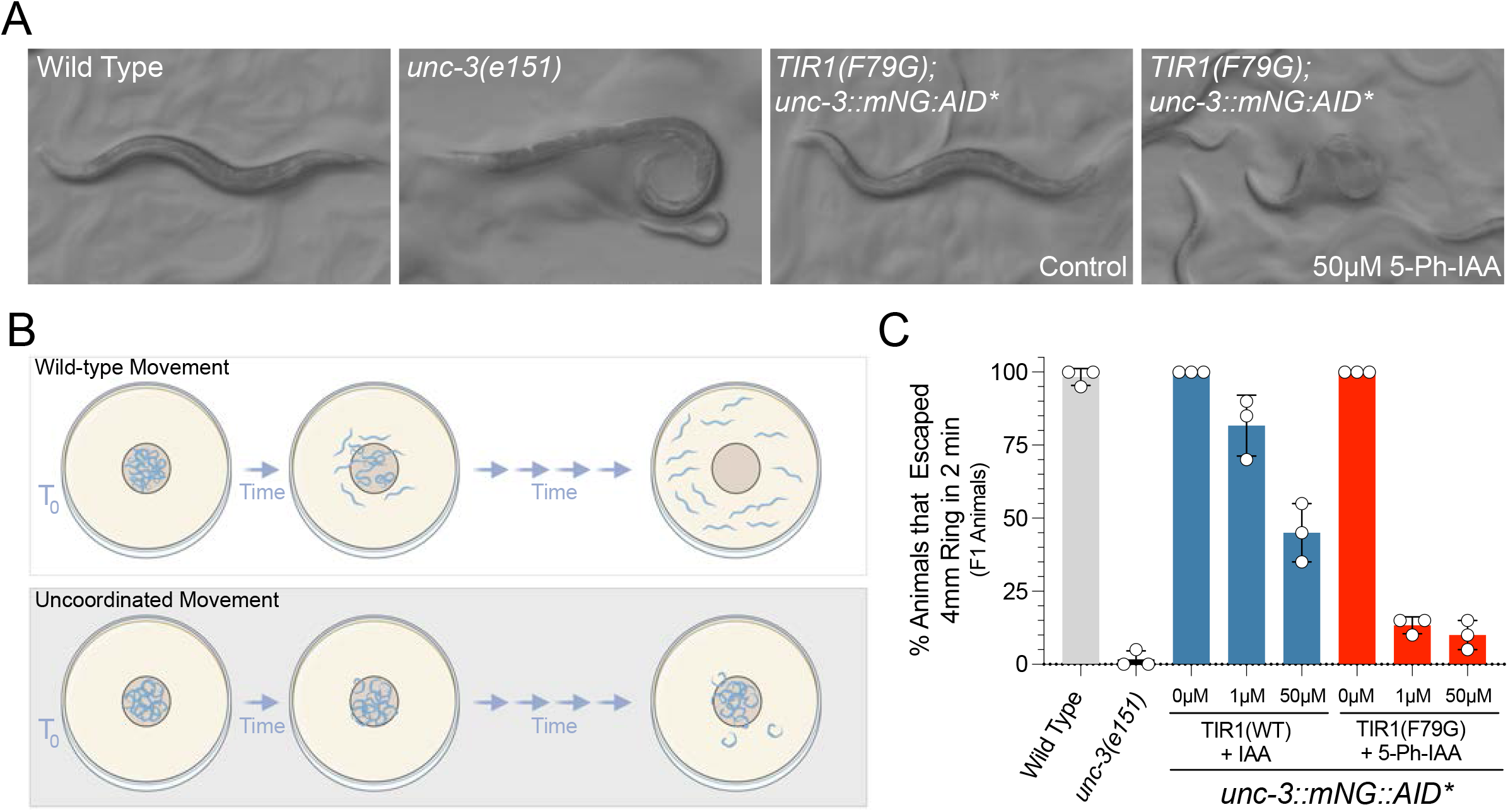
The *C*.*e*AIDv2 system generates more penetrant uncoordinated phenotypes than the _*At*_TIR1(WT)/IAA pairing for an AID-tagged allele of UNC-3. **(A)** The plate phenotypes of wild-type, *unc-3(e151)*, and _*At*_*TIR1(F79G); unc-3::GFP::AID* animals ± 5-Ph-IAA. Loss-of-functional alleles of *unc-3* cause animals to exhibit the “coiler” phenotype in which the tails of animals are paralyzed and coiled. **(B)** Depiction of the modified motility assay in which animals are placed in the center of a defined region of a solid media plate and allowed to freely move for a defined period of time. Animals were scored as uncoordinated (*unc*) if they failed to move out of the prescribed circle. **(C)** Quantification of the animals in the modified mobility assay. Each dot represents the data from four separate experiments containing 5 animals per circle.

Genes encoding essential, ubiquitously expressed RNA binding proteins are typically difficult to study using standard genetics or RNAi because maternal contribution and partial rescue of these highly expressed genes confounds analyses, and for RNAi, the half-life of these proteins is typically long. An example of this class of gene is the *C. elegans* ortholog of HNRNPA2, *hrpa-1* (Ryan and Hart 2021; Ryan *et al*. 2021). Null mutations of *hrpa-1, hrpa-1(ok963)*, can be propagated in balanced strains and homozygous *hrpa-1(ok963)* animals segregating from these animals exhibit a dramatic reduction in brood size and the resulting F1 progeny of these animals exhibit a fully penetrant larval lethal phenotype (Ryan and Hart 2021). Homozygous animals expressing a CRISPR-engineered, AID*-tagged version of *hrpa-1, hrpa-1(zen91)*, exhibit a wild type growth phenotype indicating that the AID* tag does not interfere with normal development. Consistent with previous AID*-tagged proteins, HRPA-1::AID*::TEV::FLAG animals expressing _At_TIR1(WT) exhibit a reduction in expression of the AID*-tagged target in the absence of IAA, when compared to animals expressing the _*At*_*TIR1(F79G)* allele (Figure 7A and B). We next used the *hrpa-1::AID*::TEV::FLAG* allele to determine if the *C*.*e*.AIDv2 system was more efficient at degrading target proteins than the standard _*At*_TIR1(WT) variant. To do this, we hatched transgenic animals on solid NGM media containing either IAA or 5-Ph-IAA (at the indicated concentration) and allowed these animals to grow to a young adult stage of development. Quantitative westerns (n = 4) indicate that both TIR1 variants strongly deplete HRPA-1::AID*::TEV::FLAG. We next calculated the fold reduction of HRPA-1::AID*::TEV::FLAG expression in each case and determined that the *C*.*e*.AIDv2 system was reproducibly more effective at depleting the AID*-target protein regardless of auxin analog concentration (Figure 7C). Treatment of _*At*_*TIR1(F79G)*; *hrpa-1::AID*::TEV::FLAG* with 5-Ph-IAA lead to phenotypes that superficially resemble phenotypes exhibited by homozygous *hrpa-1(ok963)* animals (Figure 7D). To compare the TIR1 two variants at the phenotypic level, we quantified brood size and percent of arrested F1 broods. Consistent with a strong depletion of *hrpa-1::AID*::TEV::FLAG* by both TIR1 alleles, addition of auxin analogs to the growth media strongly affected brood sizes (Figure 7E). In contrast to the nonpenetrant F1 arrest phenotypes of _*At*_*TIR1(WT)* animals on IAA-containing media, treatment of _*At*_*TIR1(F79G); HRPA-1::AID*::TEV::FLAG* animals with 5-Ph-IAA lead to a fully penetrant F1 embryonic or larval arrest phenotype that is identical to those measured in homozygous *hrpa-1(ok963)* animals (Figure 7F). We conclude that the C.e.AIDv2 version of the auxin degradation system more efficiently depletes AID*-tagged target proteins.

**Figure 7.**
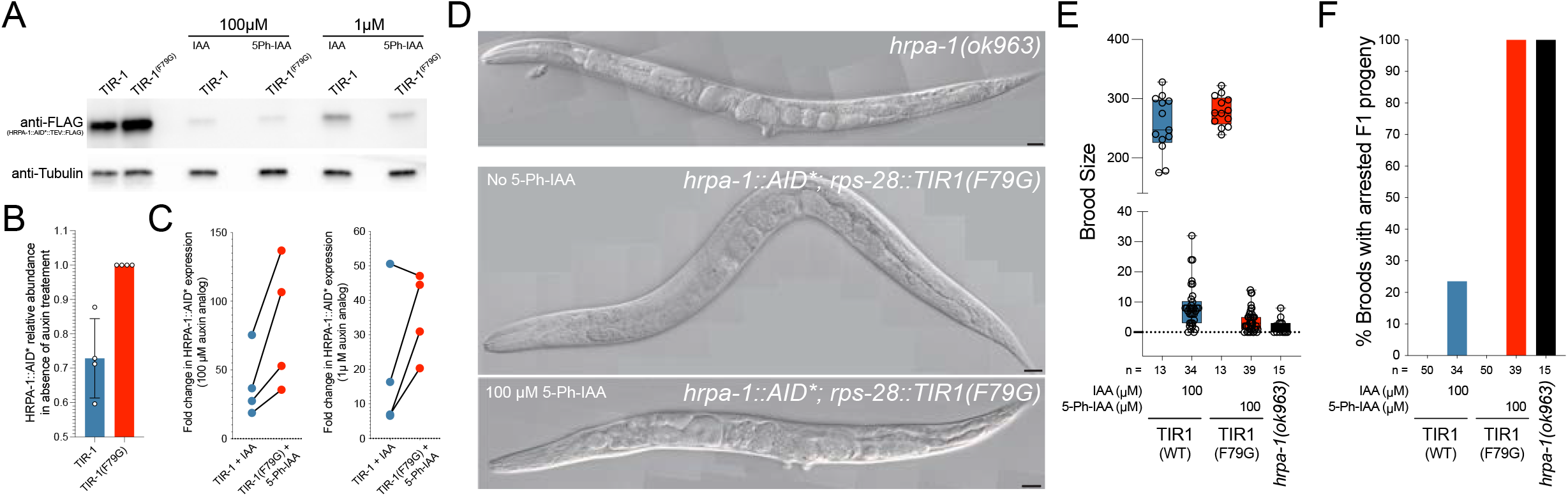
The *C*.*e*AIDv2 system phenocopies *hrpa-1(0)* phenotypes with a AID-tagged *hrpa-1* allele. **(A)** Quantitative western blots depicting the levels of HRPA-1::AID::TEV::FLAG in transgenic animals exposed to the indicated auxin analogs. For drug treatments, indicated animals were incubated from hatching to young adulthood with the indicated concentration of auxin analogs. **(B)** Quantification of the relative levels of HRPA-1::AID::TEV::FLAG in animals expressing _*At*_*TIR1(WT)* or _*At*_*TIR1(F79G)* on normal solid NGM media (n = 4). **(C)** Calculation of the relative fold reduction in HRPA-1::AID::TEV::FLAG expression in _*At*_*TIR1(WT)* or _*At*_*TIR1(F79G)* expressing animals exposed to auxin or auxin analog. **(D)** Pictomicrographs of representative young adult *hrpa-1(0)* animal or _*At*_*TIR1(F79G); hrpa-1::AID::TEV::FLAG* animals that have been grown on normal media or media containing 100µM 5-Ph-IAA as described in A and B. **(E)** Quantification of the brood size of wild-type, *hrp1-1(0)* and animals expressing *hrpa-1::AID::TEV::FLAG* and one of the two TIR1 variants on or off the indicated auxin analogs. **(F)** Calculation of the percentage of broods with arrested F1 progeny.

## DISCUSSION

The ability to rapidly and conditionally deplete target proteins using the heterologous TIR1 degradation system has dramatically improved the utility of the *C. elegans* model for dissecting aspects of multicellular development (Zhang *et al*. 2015). This is in part due to the relatively modular structure of the system which requires a single genetic modification of a target gene to insert the AID* epitope and the heterologous expression of a single E3 adaptor protein, TIR1, that, in response to the addition of an auxin analog, rapidly targets the AID*-tagged protein for degradation. This approach has enabled many gene products that were refractory to genetic manipulation to be studied in detail. In the intervening six years from its first description in *C. elegans*, the number of AID-tagged genes that have already been generated is estimated to be in the hundreds with more than 20 currently available at the *C. elegans* Genome Stock Center (CGC). For a small number of target genes, two features of the original system still exist. First, the classical allele of TIR1 exhibits a level of activity in the absence of added auxin/IAA ligand that inappropriately reduces target gene expression. As outlined in this manuscript, this ligand-independent activity is substantial and can range from 20% (for AID*::GFP and HRPA-1::AID*) to 65% (in the case of AMA-1::AID*::GFP). Relatively mild reductions of AID*-tagged proteins may not be physiologically important for many AID* targets but for other important developmental genes, this off-target activity makes the classic AID system unusable. Second, we and others find that auxin/IAA concentrations typically used to inactivate AID* target genes in *C. elegans* induce a physiological response that may occlude or modulate phenotypic observations in as of yet, unanticipated ways (Bhoi *et al*. 2021; Loose and Ghazi 2021). Our analysis of *gcs-1pro::GFP* expression indicates that at least one of these responses can occur at levels above 0.5mM IAA and are independent of TIR1 expression.

In this manuscript, we describe a modification to the TIR1 protein that solves both of these problems. Specifically, mutation of the phenylalanine 79 to a glycine residue prevents ectopic TIR1 activity while maintaining the ability to regulate AID*-tagged gene destruction through the addition of a synthetic IAA analog, 5-Ph-IAA. Further analysis of the activity of the _*At*_TIR1(F79G) variant also indicates that it exhibits a fortuitous increase in relative activity that generates more penetrant loss-of-function phenotypes for several AID*-tagged genes than those elicited with the classical TIR1 (*unc-3* and *hrpa-1*). This enables experiments to be performed at auxin analog concentrations that do not elicit undesired phenotypic consequences. The reduction in off-target effects and increase in activity can arise from a number of biochemical features that could be altered by modifying the ligand binding pocket of TIR1. We hypothesize that the inappropriate degradation of AID*-targets that occurs without the addition of exogenously added auxin may be facilitated by auxin-related indoles, whose origin may be dependent on the *E. coli* bacterial food source. It is known that commensal bacteria provide indole-related compounds to developing larva and these indoles induce phenotypic changes(Sonowal *et al*. 2017). Furthermore, we favor this hypothesis because we have noted that several sensitive AID*-tagged genes exhibit variably penetrant phenotypes that track proportionally with the age of the bacterial lawn used to culture animals. By altering the binding pocket of TIR1, the TIR1(F79G) variant may no longer be able to bind these food-derived indoles. The TIR1(F79G) variant would then be activated exclusively by the engineered 5-Ph-IAA ligand leading to enhanced experimental efficacy and tractability.

An important feature of the C.e.AIDv2 system described here is that previously existing AID*-tagged genes remain targetable with this system. This feature can be exploited in two ways. Using the CRISPR guide and repair template we used to generate the initial F79G variant, any other single copy TIR1(WT) transgene derived from the original TIR1 sequences described in Zhang et al. can be easily engineered to express the 5-Ph-IAA inducible variant, including the recent array of tissue specific drivers described in Ashley et al 2001 (Figure 2A)(Zhang *et al*. 2015; Ashley *et al*. 2021; Vo *et al*. 2021). Alternatively, we have engineered the ubiquitously expressed *rps-28pro::TIR1(F79G)* allele to contain unique, CRISPR targetable sites that flank the *rps-28* promoter sequences that can be utilized for genomic engineering and promoter preplacement. In this case, alternative promoters can easily be exchanged by homology-directed repair (HDR) (Figure S3). This method has the added advantage in that the TIR1(F79G) allele expressed from the single copy *cshIs140* allele also expresses a highly visible mCherry::HIS-11 reporter, whose ubiquitous expression will change to the expression pattern programed by any recombinant promoter sequences.

Finally, this engineered system is ripe for further enhancement at the molecular/genetic and chemical levels. In other heterologous systems, TIR1 has been modified to contain a nuclear localization sequence that targets TIR1 activity to the nucleus. This modification increases the ability of TIR1 to degrade several, high abundance nuclear proteins (Kanke *et al*. 2011). Fusion of a Skp1 ortholog, a component of the SCF complex, to TIR1 also improves degradation kinetics and loss-of-function phenotypes of AID*-tagged target genes that are refractory to TIR1 alone suggesting that these additions may also improve the TIR1(F79G) activity in *C. elegans*. Finally, 5-Ph-IAA does not appear to penetrate the eggshell. Previously described chemical modifications of IAA increase have been demonstrated to alleviate this limitation, enabling more sophisticated temporal studies of developmental processes to be addressed. It seems likely that similar modifications to 5-Ph-IAA may improve the activity of the *C*.*e*.AIDv2 system.

## ACKNOWLEDGEMENTS

We would like to thank members of the Hammell and Matus laboratories for critical review of this manuscript. We received strains from Oliver Hobert and from the Caenorhabditis Genetics Center (CGC), which is funded by NIH Office of Research Infrastructure Programs (P40 OD010440).

## FUNDING

DQM is funded by the NIH NIGMS (R01GM121597) and the Damon Runyon Cancer Research Foundation (DRR-47–17). AYZ is supported by the NIH NIGMS (R35GM124828). MAQM was supported by an NIH NIGMS Diversity Supplement (R01GM121597) and is currently funded by the NIH NCI (F30CA257383). TNM-K is supported by the NIH NICHD (F31HD100091). FEQM is supported by an NIH NIGMS Diversity Supplement (R01GM121597). JDW is supported by the NIH NIGMS (R01GM138701). SE and AM were supported by NIH NIGMS R35 GM130311. Cold Spring Harbor Laboratory, the Rita Allen Foundation, and NIH NIGMS (R01GM117406) support CMH.

## CONFLICTS OF INTEREST

None declared.

## Supplemental Figures

**Figure S1.**
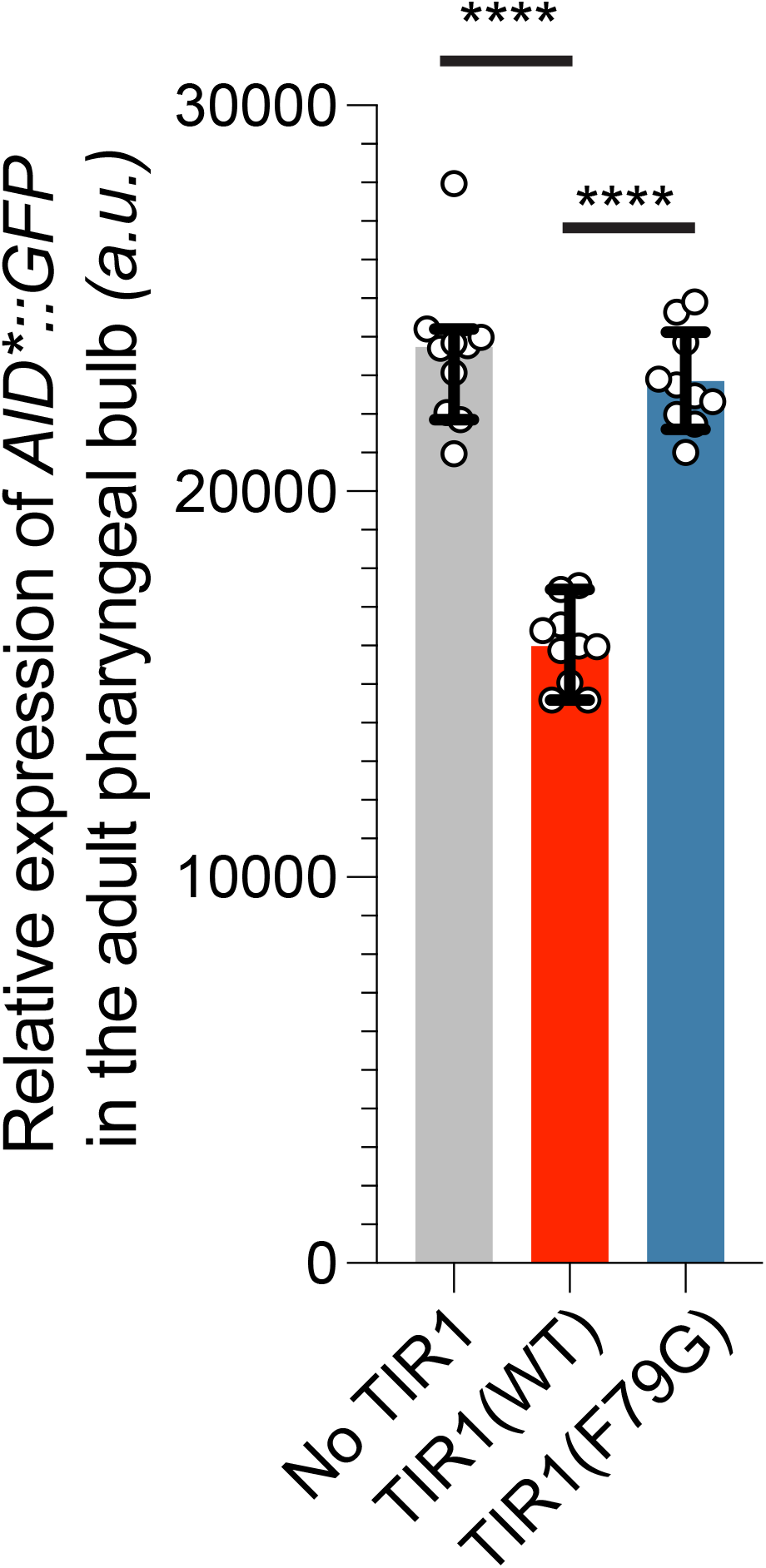
The relative expression levels of AID*::GFP (*ieSi58*) in animals expressing TIR1 variants. The relative expression levels of the AID*::GFP reporter were quantified by measuring the pixel volumes of identically sized regions of the terminal pharyngal bulbs of young adult-staged animals. Data presented as the median with SD (n = 10 animals examined for each TIR1 transgene, and **** = P < 0.0001 by a Mann Whitney U-test).

**Figure S2.**
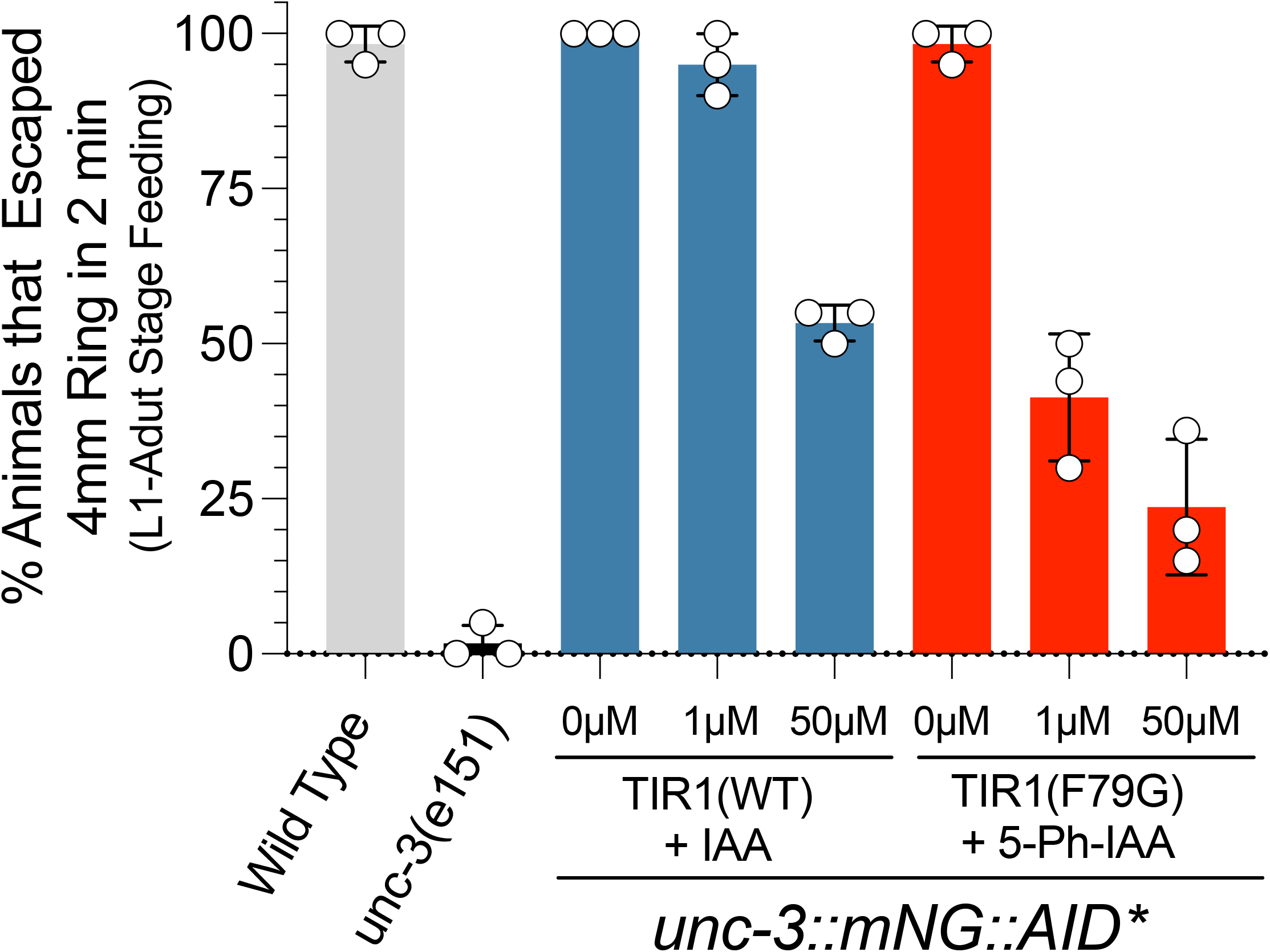
The *C*.*e*AIDv2 system generates more penetrant uncoordinated phenotypes than the _*At*_TIR1(WT)/IAA pairing for an AID-tagged allele of *unc-3* when animals are exposed to 5-Ph-IAA during pos-embryonic development. Quantification of the indicated animals in the modified mobility assay that have been exposed to 5-Ph-IAA from hatching to early adulthood. Each dot represents the data from four separate experiments containing 5 animals per circle.

**Figure S3.**
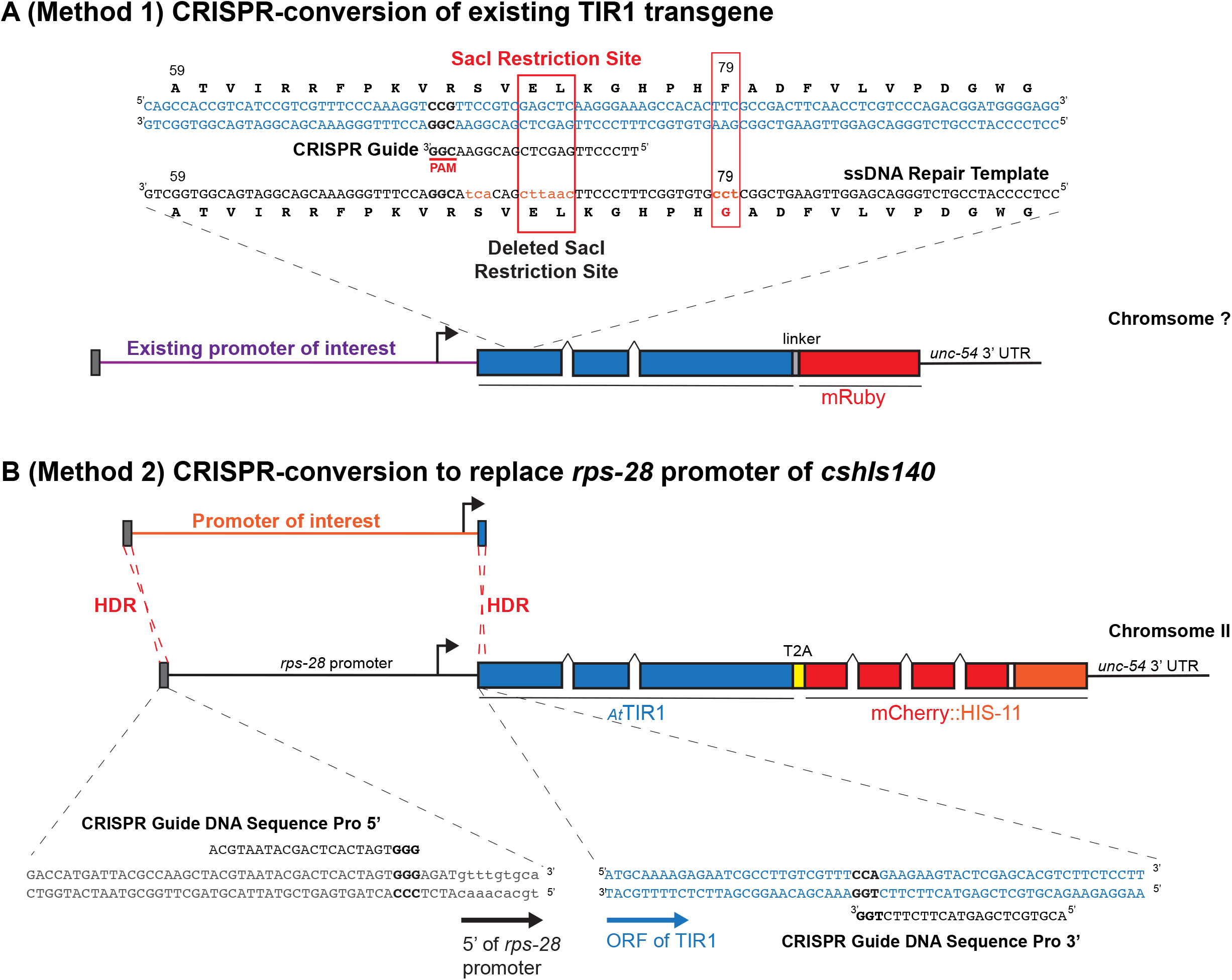
Two CRISPR-mediated methodologies to adapt the *C*.*e*.AIDv2 system. **(A)** For existing TIR1 drivers based on the original Zhang et al. TIR1 coding sequences, a single sgRNA guide/ssDNA oligo conversion strategy can be employed to generate the TIR1(F79G) variant. Using the sg RNA and repair template outlined in the methods and oligos section, conversion of previous TIR1 alleles results in the deletion of a SacI restriction site. Therefore, when successful editing has occurred, a PCR product that spans this region will lack the SacI restriction site and can be used to identify successful edits by PCR. **(B)** Alternatively, two CRISPR guides can be used to remove the *rps-28* promoter and HDR can be used to recombine in the promoter of interest.

---

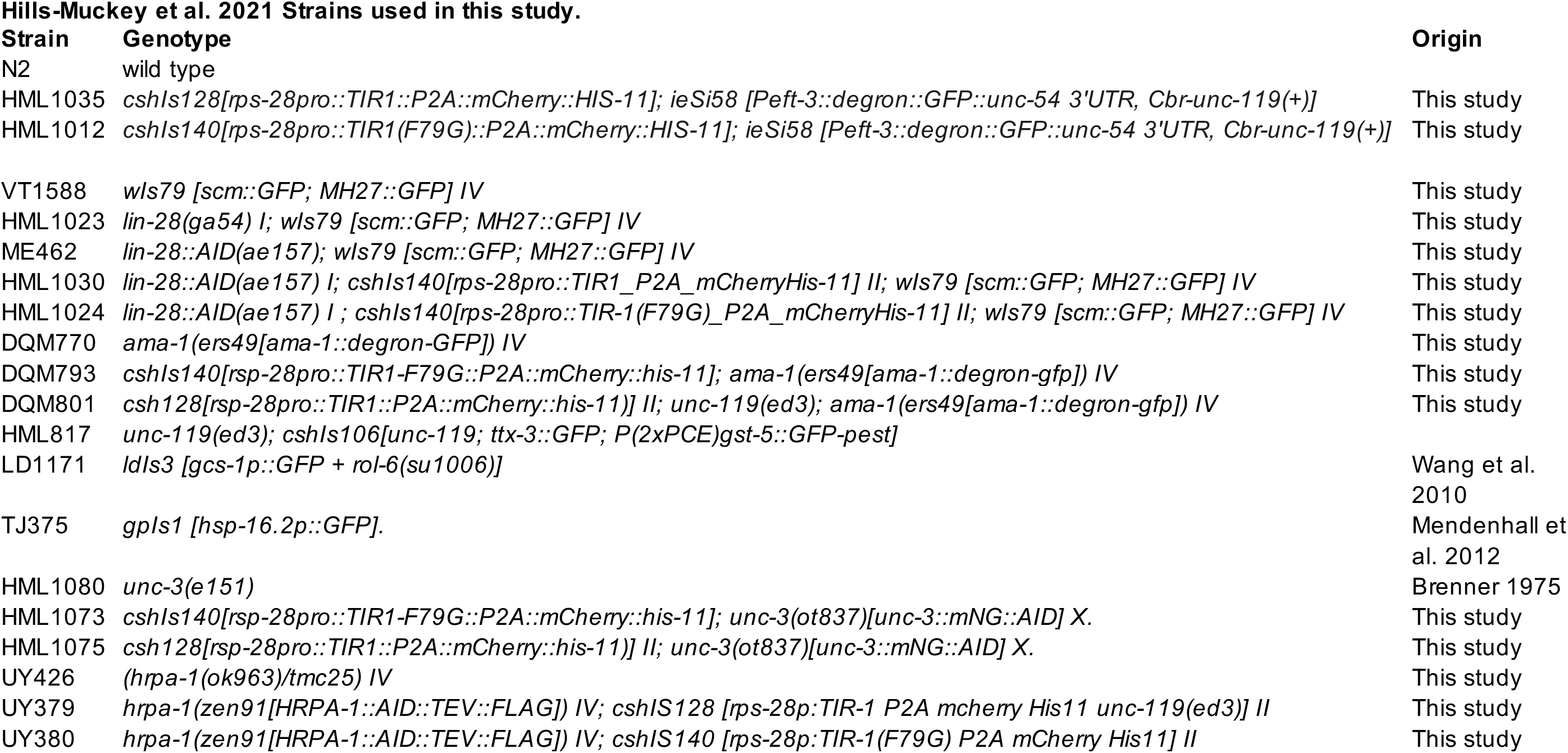

## Notes

### Competing Interest Statement

The authors have declared no competing interest.

### Summary of Updates

Spelling error of one of the authors.

